# AptaDiff: de novo design and optimization of aptamers based on diffusion models

**DOI:** 10.1101/2023.11.25.568693

**Authors:** Zhen Wang, Ziqi Liu, Wei Zhang, Yanjun Li, Yizhen Feng, Shaokang Lv, Han Diao, Zhaofeng Luo, Pengju Yan, Min He, Xiaolin Li

## Abstract

Aptamers are single-stranded nucleic acid ligands, featuring high affinity and specificity to target molecules. Traditionally they are identified from large DNA/RNA libraries using in vitro methods, like Systematic Evolution of Ligands by Exponential Enrichment (SELEX). However, these libraries capture only a small fraction of theoretical sequence space, and various aptamer candidates are constrained by actual sequencing capabilities from the experiment. Addressing this, we proposed AptaDiff, the first in silico aptamer design and optimization method based on the diffusion model. Our Aptadiff can generate aptamers beyond the constraints of high-throughput sequencing data, leveraging motif-dependent latent embeddings from variational autoencoder, and can optimize aptamers by affinity-guided aptamer generation according to Bayesian optimization. Comparative evaluations revealed AptaDiff’s superiority over existing aptamer generation methods in terms of quality and fidelity across four high-throughput screening data targeting distinct proteins. Moreover, Surface Plasmon Resonance (SPR) experiments were conducted to validate the binding affinity of aptamers generated through Bayesian optimization for two target proteins. The results unveiled a significant boost of 87.9% and 60.2% in RU values, along with a 3.6-fold and 2.4-fold decrease in KD values for the respective target proteins. Notably, the optimized aptamers demonstrated superior binding affinity compared to top experimental candidates selected through SELEX, underscoring the promising outcomes of our AptaDiff in accelerating the discovery of superior aptamers.

**Key Points:**

- We proposed AptaDiff, the first in silico aptamer design method based on the diffusion model. Aptadiff can generate aptamers beyond the constraints of high-throughput sequencing data.
- Aptadiff can optimize aptamers through affinity-guided generation via Bayesian optimization within a motif-dependent latent space, and the affinity of the optimized aptamers to the target protein is better than the best experimental candidate from traditional SELEX screening.
- Aptadiff consistently outperforms the current state-of-the-art method in terms of quality and fidelity across high-throughput screening data targeting distinct proteins.

## Introduction

Aptamers, single-stranded oligonucleotides, exhibit high affinity and specificity for specific targets due to their three-dimensional folding structure [1, 2, 3]. Like antibodies, aptamers have a wide range of applications, including targeted therapy [4, 5, 6], diagnostics [7, 8], biosensor technology [9] and biomarker discovery [10]. Aptamers offer several advantages over traditional protein antibodies. First, they possess a broad target range, encompassing metal ions, nucleotides, amino acids, growth factors, proteins, viruses, bacteria, living cells, and tissue samples [11, 12, 13, 14, 15, 16, 17]. This versatility enhances their applicability across various disciplines. Additionally, aptamers exhibit low immunogenicity and exceptional tissue penetration due to their oligonucleotide nature. Their small size and low molecular weight enable efficient tissue penetration and rapid in vivo target localization, surpassing antibodies [18, 19]. Moreover, aptamers are cost-effective and easily synthesized in vitro using DNA synthesis, PCR, or RNA transcription, ensuring minimal batch-to-batch variations and excellent repeatability. Lastly, aptamers demonstrate thermal stability, as their denaturation process is reversible, allowing them to recover their three-dimensional structure after exposure to high temperatures [20].

Aptamers are identified by the systematic evolution of ligands by exponential enrichment (SELEX). SELEX involves iterative cycles of affinity-based selection and amplification, wherein aptamer candidates demonstrating robust binding affinity to the target are successively enriched and isolated in each round. This iterative process ultimately results in the acquisition of aptamers with pronounced binding capacity for specific targets [21, 22]. In recent years, with the advancement of high-throughput sequencing technologies, it has become possible to screen a vast number of nucleic acid aptamer candidates using High-Throughput SELEX (HT-SELEX) techniques. However, compared to the vast potential sequence space, current findings suggest that aptamers with desired properties for specific tasks are exceptionally rare [23]. The theoretical sequence space of a DNA aptamer library containing a 40-nt random region is on the order of 10^25^ (~ 4^40^), whereas the size of manually prepared SELEX input libraries currently stands at around 10^15^, with sequencing throughput of approximately 10^6^. Consequently, we can only assess a very small fraction of the theoretical diversity, making it extremely challenging to select aptamers that specifically bind to target proteins within this immense sequence space. Therefore, efficient computational methods for handling high-throughput sequencing data are crucial in aptamer development.

To effectively identify high-quality aptamers from high-throughput sequencing data, Singh et al. [24] used a three-dimensional non-fouling porous hydrogel with immobilized target proteins for high-affinity aptamer selection. In addition, some machine learning and deep learning models have been employed for aptamer assessment. AptaNet utilizes a multilayer perceptron (MLP) trained on an aptamer database to predict whether an aptamer binds to a protein [25]. Rube et al. [26] used interpretable machine learning to predict protein-ligand binding affinities from sequencing data. Ali Bashir et al. [27] partitioned aptamer sub-libraries by affinity level using Particle Display (PD) and trained machine learning models on these data to predict high-affinity DNA aptamers from experimental candidates. While these methods are valuable for analyzing existing sequences, they are limited to exploring the sequences that actually exist in the database. Simulation-based methods have been reported for sequence generation [28, 29, 30], but these methods require prior knowledge of sequence motifs. Thus, motif mining plays a crucial role in identifying protein-binding sites on DNA or RNA, aiding in understanding gene regulation and control mechanisms. Several machine learning-based methods have been developed for motif detection [31, 32, 33]. In addition to simulation-based methods, some approaches utilize experimental mutation information (nucleotide substitutions) for aptamer candidate discovery [29, 27]. While mutation information is advantageous for aptamer discovery, this approach comes with considerable uncertainty.

We focused on aptamer generation methods based on deep learning. Jinho IM et al. [34] employ LSTM networks to generate aptamer sequences similar to the training data by learning from high-throughput experimental data. Furthermore, other neural network-driven generative models like Generative Adversarial Networks (GANs) [35] and Variational Autoencoders (VAEs) [36] have been proposed for nucleic acid sequence generation [37, 38, 39]. Recently, denoising diffusion probability models (DDPMs) [40, 41, 42] have achieved significant successes in computer vision field for their superior generative capabilities and have been applied to domains like small molecule and protein design [43, 44, 45, 46, 47, 48]. Diffusion models operate by continually introducing Gaussian noise to perturb training data and subsequently learning to reverse the noise process for data recovery. After training, diffusion models can incorporate randomly sampled noise into the model and generate data by learning the denoising process.

In this paper, we introduce AptaDiff, a conditional discrete diffusion model for aptamer generation. It is the first computational model based on diffusion generative models in the field of aptamer generation. AptaDiff shares similarities with the multinomial diffusion [49] for categorical data but has been tailored specifically for aptamers. We enhance the aptamer sequence generative process by extending the diffusion model with motif-dependent latent representations as conditions (see Fig. 1). The latent space is obtained through a Variational Autoencoder (see Fig. 1b), where sequences form clusters based on motif structures (see Fig. 1d). This operation effectively equips the diffusion process with a latent space, enhancing the model’s interpretability. Utilizing latent representations, we generated aptamers absent from high-throughput sequencing data and optimized them for greater affinity to target protein through affinity-guided Bayesian optimization (see Fig. 1f).

**Fig. 1.**
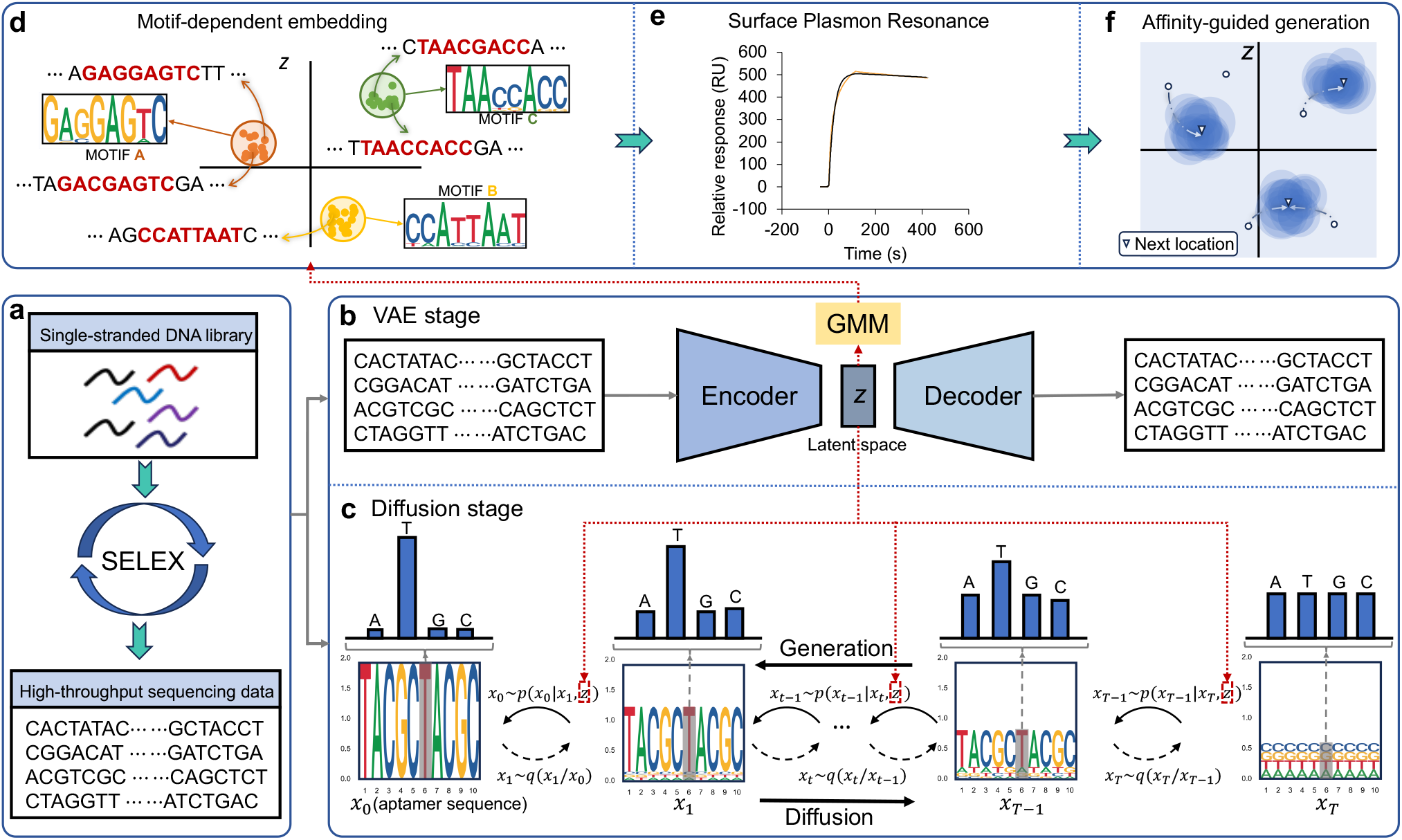
Schematic view of the AptaDiff architecture, with **(a), (b)**, and **(c)** representing the training process of AptaDiff. **(a)** Training data comes from high-throughput sequencing data from SELEX experiments. AptaDiff comprises a VAE phase and a diffusion phase. **(b)** In the VAE phase, a latent space with motif-dependent information can be obtained by a specific VAE (the latent space is shown in **z** in this figure). **(c)** During the diffusion stage, we model the training data using a Multinomial Diffusion conditioned on the motif-dependent embedding. The data used for diffusion is the aptamer sequence. In subfigure **c**, the diffusion process of the aptamer sequences is displayed below, while the diffusion state of individual nucleotides within the sequences is shown above. **(d)** The VAE constructs the latent space based on sequence similarity. **(e)** The binding affinity between the aptamer and the target was measured by surface plasmon resonance experiment. **(f)** AptaDiff can propose candidate aptamers through Bayesian optimization based on the binding affinity distribution by converting the latent representation into a probabilistic model.

We demonstrate that the generated sequences exhibit characteristics similar to real HT-SELEX sequences across various metrics, and the optimized sequences show stronger affinity to the target protein. We believe that AptaDiff represents an effective technique for designing aptamer sequences as candidate therapeutics and opens up a promising new avenue for aptamer-based therapies.

## Materials and Methods

### Preliminaries

#### Variational Autoencoders

Variational autoencoders (VAE) comprise an encoder neural network and a decoder neural network, where the encoder ℰ_*ϕ*_ transforms input sequences (x) into a latent distribution (q_*ϕ*_(z∣x)), and the decoder 𝒟_*θ*_ learns to reconstruct input data from the latent representation (z) by learning (*p*_*θ*_ (x∣z)), where *ϕ* and *θ* represent model parameters. The whole framework can be trained by minimizing the reconstruction objective *d*(𝒟 (ℰ (x), x)). As a generative model, VAE’s evaluation hinges on model evidence, yet computing the model evidence (*p*_*θ*_ (X)) for a dataset 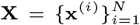 proves computationally challenging. Alternatively, we turn to maximizing the Evidence Lower Bound (ELBO), denoted as ℒ_*VAE*_(*θ, ϕ*;X), to gauge the model’s description of the dataset using Jensen’s inequality. The whole VAE can be effectively optimized by:

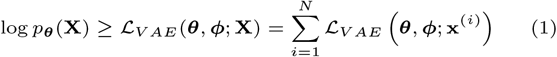

where

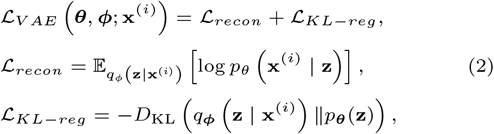

which is a reconstruction loss ℒ_*recon*_ combined with a regularization term ℒ_*KL−reg*_. *D*_KL_ (*p*∥*q*) stands for the Kullback–Leibler divergence between distributions *p* and *q*. The reconstruction loss in practice is calculated as *L*_2_ norm or cross-entropy loss. In this paper, we refer to the negative value of ELBO as the model loss.

#### Diffusion Probabilistic Models (DPMs)

Diffusion probabilistic models (DPMs) are a class of generative models based on a reversible, discrete-time diffusion process. Given data ***x***_0_, a diffusion probabilistic model defines two Markov chains of diffusion processes. The forward diffusion process consists of predefined variational distributions *q*(***x***_*t*_|***x***_*t−*1_) that gradually reduce the signal-to-noise ratio over time steps *t* ∈ {1, …, *T*} until the data distribution approximately reaches the prior distribution. The generative diffusion process consists of learnable distributions *p*(***x***_*t−*1_|***x***_*t*_) that start from the prior distribution and transform it to the desired distribution by progressively denoising noisy data.

#### Dataset

The training data used in this study comprises 4 independent HT-SELEX datasets, denoted as datasets A, B, C, and D. These datasets are directed toward four distinct target proteins. Specifically, datasets A and B were derived from HT-SELEX experiments conducted by ourselves (see Supplementary Section 2 for details), while datasets C and D were obtained from two public SELEX datasets provided by DNA Data Bank of Japan (DDBJ) [50, 51]. The original HT-SELEX data consists of variable random regions flanked by fixed primer regions. The target sequence lengths in datasets A, B, C, and D are 36 nt, 36 nt, 30 nt, and 40 nt, respectively. For detailed information about each dataset and their corresponding target proteins, please refer to Supplementary Section 2 and Table S9 for specific references and descriptions.

#### Overview of AptaDiff

Here, we propose AptaDiff, a novel conditioning diffusion-based generative model, to generate new aptamer sequences that are not included in the input SELEX dataset. The AptaDiff network comprises two main stages, as shown in Fig. 1. In the first stage, a latent space with motif-dependent information can be obtained by a specific VAE (see Fig. 1b). In the second stage, we can then model the training data using a Multinomial Diffusion [49] conditioned on the motif-dependent embedding (see Fig. 1c). This design effectively equips the diffusion process with a low-dimensional latent space. Using motif-dependent embedding as the condition can guide the generation of aptamers, thereby resulting in better generative performance than existing models.

In the proposed model, the initial stage is to train a VAE to learn the low-dimensional motif-dependent aptamer representation (Fig. 1d). Notably, motifs are neither explicitly generated by the model nor identified through direct comparison of aptamers. Instead, motifs are implicitly learned through a specific VAE. Following Iwano et al. [39], we constructed the VAE with a 1-D CNN-based encoder and a profile HMM decoder. Training this VAE yields a latent space with motif-dependency (a conclusion substantiated in reference [39]). The characteristics of motifs within this space are primarily captured by the profile HMM decoder. The encoder captures sequential motifs that are aligned in certain positions. Within this latent space, aptamer sequences can form clusters based on their motif structures, even though the specific details of these motifs remain unknown and implicit, as illustrated in Fig. 4a.

#### Details of the Diffusion Stage

Our approach to the aptamer sequence design problem builds on Multinomial Diffusion, a likelihood-based model for categorical data [49]. We follow the conventions and notation set by Hoogeboom et al. [49], which we review here.

The forward diffusion process for nucleotide (base) types is defined as follows:

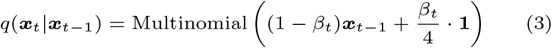

where ***x***_*t*_ will be represented in one-hot encoded format ***x***_*t*_ ∈ {0, 1}^*K*^ (*K* = 4). Specifically, for category *i, x*_*i*_ = 1 and *x*_*j*_ = 0 for *i* ≠ *j*. **1** is an all-one matrix. *β*_*t*_ is the probability of resampling another nucleotide over 4 types uniformly. When *t* → *T, β*_*t*_ is set close to 1 and the distribution is closer to the uniform distribution. The following probability density provides an efficient way to perturb x_0_ for timestep t during training [49].

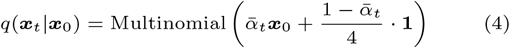

where *α*_*t*_ = 1 − *β*_*t*_ and 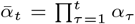. Intuitively, at each next time step, a small amount of uniform noise *β*_*t*_ is introduced over the 4 nucleotide classes, and there is a high probability (1 − *β*_*t*_) that the previous value ***x***_*t−*1_ is sampled.

The posterior *q*(***x***_*t−*1_|***x***_*t*_, ***x***_0_) can be derived from Eq.3 and Eq.4, that is:

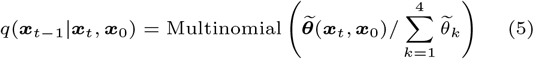

where 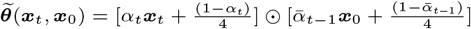.

The generative diffusion process needs to approximate the posterior *q*(***x***_*t−*1_|***x***_*t*_, ***x***_0_) to denoise. Specifically, we predict the probability vector of 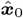 from ***x***_*t*_ and the latent embedding **z** corresponding to ***x***_0_, and then parameterize *p*(***x***_*t−*1_|***x***_*t*_) with the probability vector of 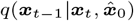. The generative diffusion process is defined as :

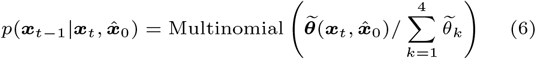

where 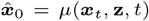 represents a neural network model, specifically utilizing the Transformer architecture [52]. This model takes the latent embedding **z** corresponding to ***x***_0_ and the aptamer state from the previous step as inputs, predicting the probability of nucleotide types at each position in the current state of the aptamer sequence. It’s important to note that, unlike the forward diffusion process, the generative diffusion process relies on both the latent embedding **z** and the aptamer state from the previous step. The key distinction between these two processes lies in the fact that the forward diffusion process introduces noise to the data, rendering it independent of the data or latent embedding **z**. On the other hand, the generative diffusion process is contingent upon the specified condition (**z**) and full observation of the previous step.

The objective of training the generative diffusion process for aptamer sequence is to minimize the expected KL divergence between Eq.6 and posterior distribution Eq.5:

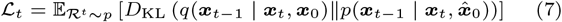

#### Handling the Diffusion Model Stochasticity

The proposed AptaDiff model consists of two types of latent representations: the low-dimensional VAE latent code **z** and the DDPM intermediate representations ***x***_1:*T*_ associated with the DDPM reverse process (which are of the same size as the input sequence ***x***_0_). The stochasticity in the generative diffusion process during the second stage of AptaDiff may lead to uncertainty in generated samples. To ensure that samples generated by AptaDiff are deterministic (i.e., controlled only by **z**), we followed [53] to share all stochasticity during the reverse diffusion process across all generated samples. Since initializing the second stage Multinomial Diffusion in AptaDiff with different random noise during sampling might be interpreted as imparting different styles to samples, sharing this stochasticity in Multinomial Diffusion sampling across samples means using the same style for all generated samples. Through this straightforward technique, we can now achieve controlled generation using the low-dimensional VAE latent code, thereby enabling deterministic sampling (see Supplementary Section 4 for details).

#### Experimental Setup

In our study, we utilized one-hot encoding to encode the data, with each aptamer having dimensions of (*L*, 4), where *L* represents the length of the aptamer, as mentioned earlier, and 4 corresponds to the length of the one-hot vector. During the training process, the batch size was set to 64. The one-hot encoding for aptamer nucleotides was as follows: Adenine (A): (1, 0, 0, 0), Cytosine (C): (0, 1, 0, 0), Guanine (G): (0, 0, 1, 0), Thymine (T): (0, 0, 0, 1).

The model was implemented using the PyTorch [54] library and Python. All sequences in the training set are carefully filtered. Sequences meeting the following three criteria are retained: (1) exact matching with primers (including forward and reverse primers), (2) precise alignment with the sequence design length, and (3) sequences read more than once. The sequences are split into training and test datasets in a 9 : 1 ratio. The model with the smallest test loss is selected through iterations. We conducted the training process using a single RTX 3090 GPU. The number of diffusion steps *T* was set to 1000. Adam was used as the training optimizer and all networks were trained up to 1000 epochs with early stopping when the test loss was not updated for 50 epochs.

#### Surface Plasmon Resonance Assay

After AptaDiff generates the sequence, we connect the obtained sequence to its fixed primer sequence, prepare it as a solid powder, and evaluate its binding affinity to the target protein through surface plasmon resonance (SPR) experiments. The instrument used for the SPR experiments (Fig. S3) was Biacore 8K (GE Healthcare). We mainly conducted SPR analysis on datasets A and B. The target proteins of datasets A and B were insulin-like growth factor-binding protein 3 (IGFBP3, UniProt: p17936) and Inactive tyrosine-protein kinase 7 (PTK7, UniProt: Q13308), respectively. Aptamers were prepared with fixed primer regions at both ends and a random region in the middle as follows: 5’-TCCAGCACTCCACGCATAAC-(36 nt random region)-GTTATGCGTGCTACCGTGAA-3’ for dataset A and 5’-ATTGGCACTCCACGCATAGG-(36 nt random region)- CCTATGCGTGCTACCGTGAA-3’ for dataset B. Aptamers screened by SELEX wet assay were used as positive controls. All evaluated sequences are listed in Supplementary Section 1 and Supplementary Table S1, S2, S4, S5, S6, S7 and S8). Aptamers are synthesized in vitro based on sequence information and prepared as a solid powder. The running buffer is 1xDPBS (1XDulbecco’s Phosphate Buffered, PH 7.4-7.6) containing 5 mM magnesium ions (*Mg*^2+^). The sensor chip is Biacore series S sensor chip CM5 (BR-1005-30, Cytiva), its gold film surface is covalently linked to the carboxymethyl dextran matrix, and the protein is covalently coupled to the carboxyl group on the surface of the sensor chip through the amino group (Fig. S3). Coupling the protein to the second channel of the CM5 chip as the experimental group, the specific steps are as follows:

- Mix equal volumes (1 : 1) of EDC (N-(3-Dimethylaminopropyl)- N’-ethylcarbodiimide hydrochloride, 0.4 M aqueous solution) and NHS (N-Hydroxy Succinyl, 0.1 M aqueous solution) to activate the carboxyl groups on the chip. The instrument parameters are injection time 420 s, and flow rate 10 *µ*L/min.
- Dilute the proteins of dataset A and dataset B with 10 mM acetic acid-sodium acetate buffer solution at pH 4.5 for covalent coupling with the carboxyl groups on the surface of the chip. The instrument parameters are a flow rate of 10 *µ*L/min and a binding time of 900 s.
- Use ethanolamine (1 M) to block the carboxyl group of the unbound protein on the chip, the instrument parameters are flow rate 10 *µ*L/min, and blocking time 420 s.

The 9*His Peptide was coupled to the first channel of the CM5 chip as a control group, and the experimental steps were exactly the same as above. Afterward, the sequences of datasets A and B flowed through the sensor chip to measure their affinity with the protein (sequences were diluted with DPBS containing 5 mM magnesium ions at a concentration of 500 nM, the flow rate was 30 *µ*L/min, and the binding time was 120 s, the dissociation time of A is 120 s, and the dissociation time of B is 300 s). In order to regenerate the sensor chip and completely remove the bound aptamers, a regeneration solution is required. The regeneration solution of protein A is 1.5 M sodium chloride with 5 mM NaoH (flow rate 30 *µ*L/min, time 60 s), and the regeneration solution of protein B is 2 mM NaoH 1.5 M sodium chloride (flow rate 30 *µ*L/min, time 30 s).

## Results

### AptaDiff shows the state-of-the-art generative performance, robustness, and generalization

We hope that the AptaDiff architecture described above can learn to generate aptamer sequences similar to real SELEX data. Specifically, we aim for the generated samples to align with the various attributes of the real SELEX data used for training and validation. Currently, there is a dearth of work focused on aptamer generation, and the evaluation of aptamer generation remains an open question. In this paper, we carefully set up our evaluation pipeline to reflect generative quality. We evaluate our model from multiple perspectives to comprehensively examine the quality of aptamer generation, including the distance between the generated sequence and the real sequence, the GC content, the minimum free energy (MFE) distribution, base distribution, the FID value, and motifs cluster analysis. We also compare our model with randomly generated sequences and the current state-of-the-art aptamer generation model—RaptGen [39]. Additionally, to validate the robustness and generalizability of our model, we conducted experiments on four independent HT-SELEX datasets (data sets A, B, C, and D, respectively for 4 different target proteins).

#### Sequence Distance Evaluation

In order to assess the distance between the sets of generated sequences and random sequences in comparison to real SELEX data, we opted for the Levenshtein distance as our distance metric. The Levenshtein distance is a method for quantifying the dissimilarity between two strings by calculating the minimum number of single-character edits (insertions, deletions, or substitutions) required to transform one string into another [55]. Following the research conducted by Ozden et al. [38], we determined the distance of each generated or random sequence to the nearest sequence in the real samples and utilized these sets of distances to represent the distance distributions of these sequence collections. The distance distribution of the real dataset was obtained by computing the minimum non-zero distance between each real sample and the entire real SELEX dataset. The distance distributions depicted in Fig. 2a indicate that, across the four datasets, our AptaDiff-generated sequences exhibit nearly identical distance distributions compared to real sequences, while randomly generated samples display different distributions. Compared with the sequences generated by RaptGen, the sequences generated by our model are closer to the distance distribution of the real sequences in the three data sets A, C, and D, and the performance on data set B is equivalent to RaptGen. This suggests that our AptaDiff model has more effectively learned the contextual characteristics of aptamer sequences.

**Fig. 2.**
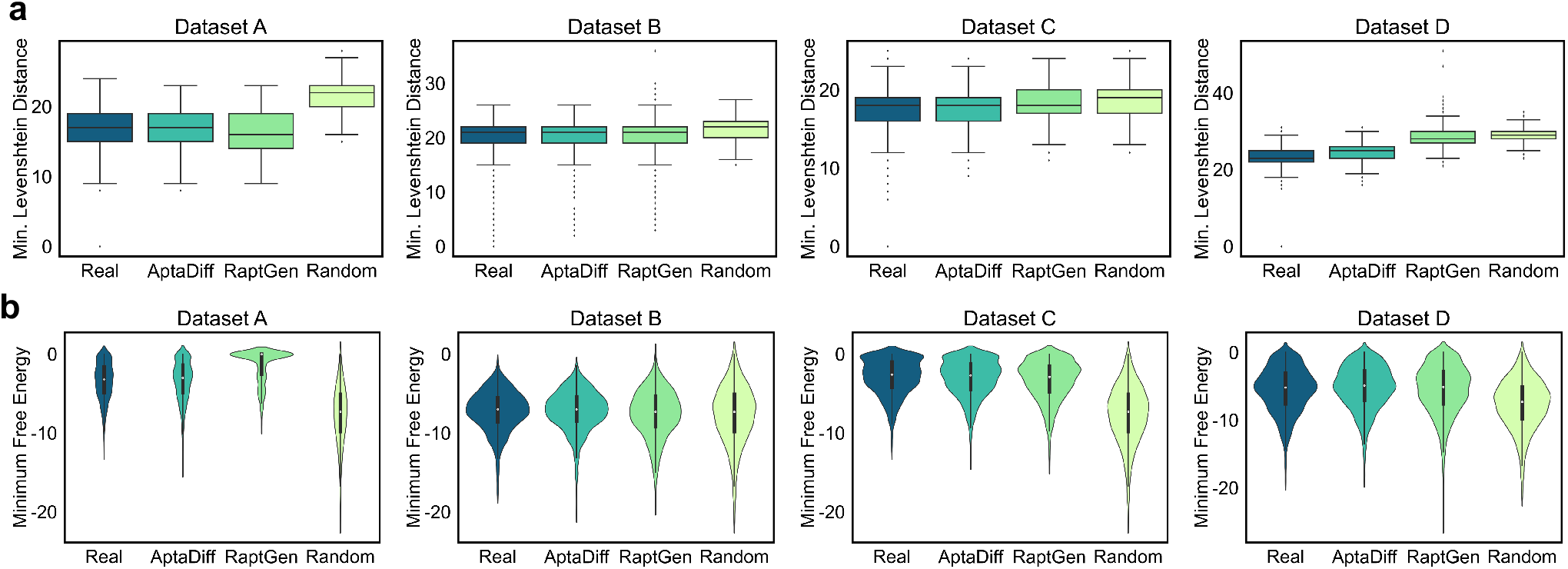
Levenshtein distance distributions (a) and Minimum Free Energy distributions (b). **(a)** The Levenshtein distances measured between the generated sequences from AptaDiff, RaptGen, and random sequences to the nearest real sequences are shown in the distributions for four datasets (A, B, C, D). To quantify the real sequences, the distance to the nearest member with a non-zero distance was considered. Both random and generated sequences have a size of 2, 000, and all real sequences used for training are included here. Box plots characterize a sample using the 25th, 50th (also known as the median or Q2), and 75th percentiles—referred to as the lower quartile (Q1), median (m or Q2), and upper quartile (Q3)—and the interquartile range (IQR = Q3-Q1), covering the central 50% of the data. Whiskers extend outside the box up to 1.5 times the IQR. Any data points outside the whiskers are considered outliers and are plotted individually as dots. The results demonstrate that, across the four datasets, the sequences generated by our AptaDiff exhibit nearly identical distance distributions compared to the real sequences. **(b)**We compared the Minimum Free Energy (MFE) distribution among 2, 000 sequences generated by AptaDiff, 2, 000 sequences generated by RaptGen, 2, 000 randomly generated sequences, and the MFE distribution of the real HT-SELEX data used for training and validation. The results indicate that, when compared to real sequences, our AptaDiff-generated sequences exhibit nearly identical MFE distributions across all four datasets. In contrast, sequences generated by RaptGen show MFE distributions that closely resemble the original sequences in some datasets, while randomly generated samples exhibit different distributions.

#### Minimum Free Energy Distribution Evaluation

The minimum free energy (MFE) of nucleic acid aptamer sequences determines the stability of their secondary structure, and as such, we aim for the MFE distribution of the generated samples to align with that of real samples. We employed the ViennaRNA package [56] to calculate the MFE of aptamer sequences. As depicted in Fig. 2b, we evaluated the MFE distribution of 2, 000 sequences each from AptaDiff, RaptGen, and randomly generated sequences against the MFE distribution of the real HT-SELEX data utilized for training and validation. The results indicate that, in comparison to real sequences, our AptaDiff-generated sequences exhibit nearly identical MFE distributions across all four datasets, while sequences from both RaptGen and random generation exhibit more distinct distributions. This suggests that AptaDiff effectively emulates real data in terms of minimum free energy.

#### GC Content and Base Distribution Evaluation

We conducted an investigation into the GC content and base distribution of the generated sequence sets. We separately computed the GC content and base distribution for 2, 000 sequences generated by AptaDiff and 2, 000 sequences generated by RaptGen. We then compared their distributions to those of 2, 000 real sequences. The GC content distribution illustrated in Fig. 3a reveals that, across the four datasets, AptaDiff-generated sequences exhibit GC content distributions more similar to real sequences compared to sequences generated by RaptGen. This similarity is particularly pronounced when the datasets have higher GC content (e.g., datasets A and B). The base distribution shown in Fig. 3b illustrates that the base distribution of the sequences generated by AptaDiff is more similar to the real sequences compared with the sequences generated by RaptGen on the 4 datasets. It is worth noting that in our analysis, we did not include the GC content and base distribution of random sequences since all four bases have an equal probability of occurrence (25%) at every position in randomly generated sequences.

**Fig. 3.**
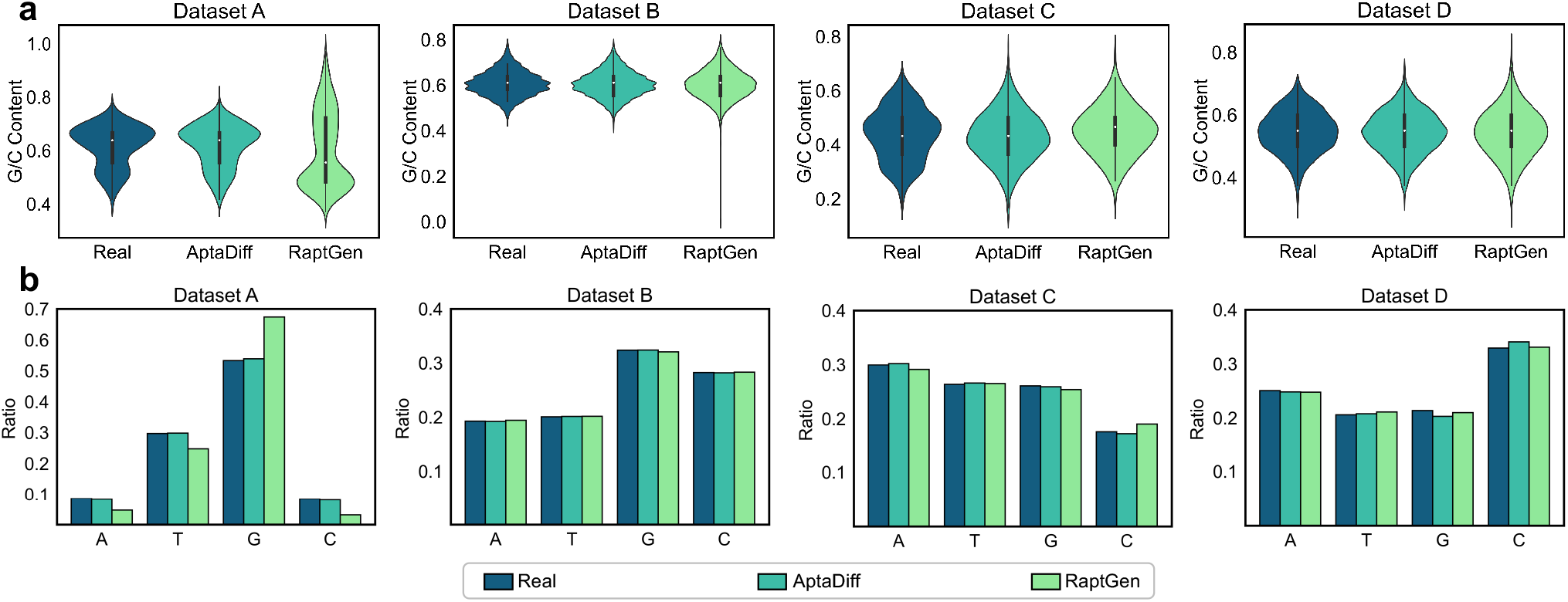
GC content distributions (a) and base distributions (b). **(a)** We conducted an analysis of the GC content distribution for the generated sequence sets. We separately computed the GC content distribution for 2, 000 sequences generated by AptaDiff and 2, 000 sequences generated by RaptGen and compared their distributions to the GC content of 2, 000 real sequences. The results demonstrate that, across the four datasets, our AptaDiff-generated sequences exhibit more similar mean values and distributions of GC content compared to the real sequences. **(b)** We conducted an analysis of the base distribution for the generated sequence sets. We separately computed the base distribution for 2, 000 sequences generated by AptaDiff and 2, 000 sequences generated by RaptGen and compared their distributions to the base distribution of 2, 000 real sequences. The results indicate that, across the four datasets, the base distribution of sequences generated by AptaDiff is more similar to the real sequences compared to sequences generated by RaptGen.

**Fig. 4.**
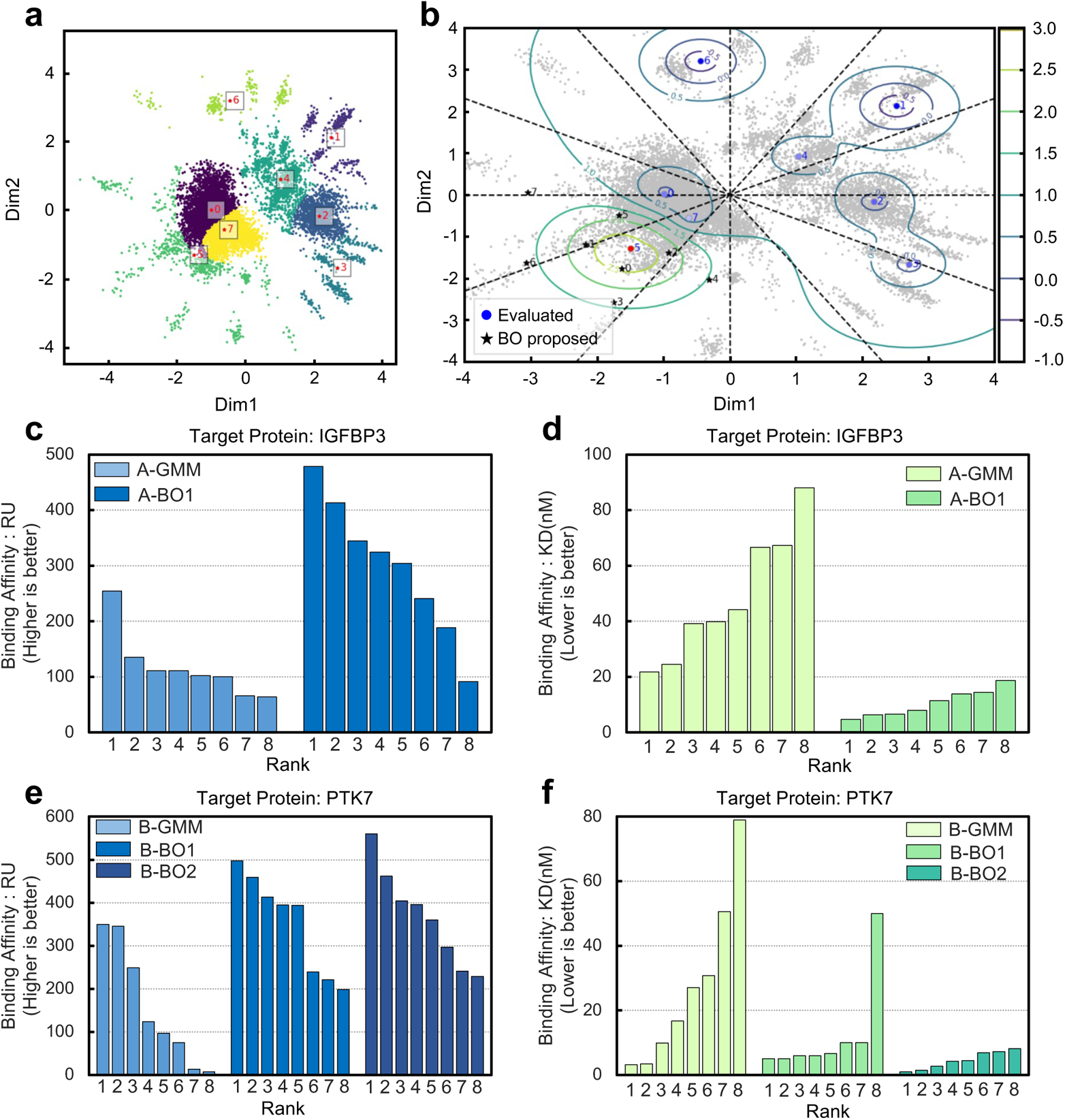
Results of Gaussian Mixture Model (GMM) and Bayesian Optimization (BO). **(a)** Visualization of the latent space for Dataset A, with clustering estimated by GMM. The colored patches represent different clusters. Initial points for Bayesian optimization are obtained from each center of the Gaussian distribution. Their IDs, positions in the latent space, and corresponding affinity values with the target protein measured by SPR experiments are shown in Supplementary Table S4. **(b)** Affinity distribution for Dataset A and proposed Bayesian optimization (BO) points. The affinity data from Supplementary Table S4 are embedded into the latent space. Gray points represent the embedded points from Fig. 4a. Contours overlaid on the embedding represent predicted affinity levels, which serve as the acquisition function for Bayesian optimization. Bayesian optimization process suggests 8 points. Circles represent the re-embedding positions of GMM centers. Red and blue colors indicate high and low binding activities, respectively. Black stars represent the positions suggested by Bayesian optimization. **(c), (d), (e), (f)**. Affinities of sequences proposed by different methods; A-GMM and B-GMM represent GMMs performed on the latent space of datasets A and B, respectively; A-BO1 and B-BO1 represent the first round of Bayesian optimization on datasets A and B, respectively; similarly, B-BO2 represents the second round of Bayesian optimization on dataset B. Rank represents the affinity ranking within the method (ranking of RU value (**c**) and KD value (**d**) of dataset A, and ranking of RU value (**e**) and KD value (**f**) of dataset B).

#### FID Evaluation

The Fréchet Inception Distance (FID) was originally introduced as a metric for evaluating the quality of images generated by generative models [57]. FID compares the distribution of generated images to that of a reference set of real images, often referred to as the “ground truth,” and has become one of the most widely used metrics for assessing the quality of generative models.

In order to compare the distribution of aptamer sequences generated by our model to real data, we adapted the FID comparison for aptamers, following these steps: (1) We randomly selected 10, 000 sequences from both the generated sequences and real sequences. (2) We extracted features from these sequences using RNA-FM [58], resulting in feature vectors of dimension (*L*, 640) for each sequence, where *L* represents the length of the aptamer sequence and 640 represents the dimension of each base embedding, thereby obtaining feature distributions for both datasets. (3) We computed the mean vectors and covariance matrices for the two distributions. (4) Finally, we calculated the Fréchet distance between the two distributions, yielding the FID score. As shown in Table 1, compared with RaptGen on 4 data sets, the FID values between the sequences generated by AptaDiff and the real sequences are smaller, indicating that the sequences we generated are more similar to the real sequences and of higher quality. Similar results were obtained by extracting features from the sequences using DNA-BERT2 [59] (see Supplementary Table S3).

**Table 1.** Quantitative Illustration of the generative performance on datasets A, B, C, and D. FID reported on 10k samples (Lower is better).

| Source | Dataset | Target Protein | FID |  |
| --- | --- | --- | --- | --- |
|  |  |  | RaptGen | AptaDiff |
| Ours | A | IGFBP3<sup>1</sup> | 7.97 | 2.51 |
|  | B | PTK7<sup>2</sup> | 6.61 | 1.67 |
| Public | C | TGM2<sup>3</sup> | 2.48 | 1.06 |
| | D | ITGαVβ3<sup>4</sup> | 2.65 | 1.58 |

- 1: IGFBP3, Insulin-like growth factor-binding protein 3. - 2: PTK7, Inactive tyrosine-protein kinase 7. - 3: TGM2, Human transglutaminase 2. - 4: ITG $\alpha$ V $\beta$ 3, Human integrin alpha V beta 3.

#### Motif Clustering Evaluation

Since HT-SELEX sequences are amplified using specific binding motifs, and motifs play a crucial role in the aptamer-target binding process, the rationality of aptamer sampling is evidently important. We aim for our designs to incorporate motifs that are similar to those present in the training data. To assess the consistency of motifs contained in samples generated by AptaDiff and RaptGen relative to real data, we employed MEME [60] as a motif clustering analysis tool to evaluate their alignment with genuine aptamers. The motif clustering results presented in Fig. S1 and S2 demonstrate that, when compared to real sequences across the four datasets, AptaDiff-generated sequences contain similar motifs. Although their performance relative to RaptGen is not significantly superior, it still reaches a considerable level, indicating that our model effectively learns the motif information of aptamer sequences.

**In summary**, our proposed method consistently generates aptamer sequences across all four datasets that align with various attributes of real SELEX data. The performance surpasses that of the current state-of-the-art aptamer generation models, demonstrating the exceptional capabilities of our model in terms of generation quality, robustness, and generalization.

### AptaDiff application in aptamer discovery for specific targets

Most existing papers in the field of biomolecular generation [61, 62, 63, 64] primarily focus on showcasing the capabilities of their models from an algorithmic design perspective but often neglect the evaluation of the effectiveness of these models in real-world drug design. Here, we verified the capability of our model in target-specific aptamer discovery through wet lab experiments. In order to provide a genuine assessment, we carefully selected two representative real-world targets (IGFBP3 and PTK7), prepared their corresponding protein crystal structures, and conducted an analysis of the binding affinities of the sequences we generated in the ensuing discussion. The original HT-SELEX data for IGFBP3 and PTK7 has 36 nt variable regions with fixed primer regions at both ends. In this study, we refer to Iwano et al. [39] to create a latent space using the variable region. According to the experiments and conclusions of Iwano et al. [39], when the dimension of the latent embedding is set to two, the VAE’s ELBO loss is small. As the embedding dimension increases, the loss tends to increase. Additionally, a lower dimension is easier to visualize and more conducive to performing Bayesian optimization. Therefore, we also adopt a two-dimensional latent space for analysis. Fig. 4a shows the generated latent embeddings.

Due to the clustering of SELEX experiment sequences in latent space based on their motifs, we, similar to Iwano et al. [39], employed a Gaussian Mixture Model (GMM) to classify the distribution and selected eight distinct points representing the latent clustering centers of the GMM. Starting from these GMM centers, we attempted to generate corresponding sequences through AptaDiff for binding activity evaluation conditionally. The obtained aptamer sequences and the coordinates of each GMM center are listed in the Supplementary Table S4 and S6. After linking the obtained sequences with their fixed primer sequences, we prepared them as solid powders and assessed their binding affinity with the target protein through SPR experiments, the principle is shown in Fig. S3. Aptamers with the highest affinity selected through SELEX screening assay (see the Supplementary Tables S1 and S2) were used as positive controls. Although more than half of the candidate sequences exhibited weak or no binding affinity, a subset of sequences such as A-GMM-5, B-GMM-0, and B-GMM-4 demonstrated notable binding affinities (the Supplementary Table S4 and S6).

The VAE model within our AptaDiff framework associates each aptamer with a low-dimensional latent vector, resulting in a compact latent space (see Fig. 4a). To predict effective candidates from the vicinity of aptamers with high binding affinity, we employed a Bayesian optimization (BO) algorithm to learn an activity distribution in the latent space (see Fig. 4b) and generated aptamers based on the embeddings recommended after optimization. For Bayesian optimization, as shown in the flowchart in Supplementary Fig. S4, we initially embedded binding affinity data (RU values^1^) obtained from GMM centers into the latent space. Subsequently, we perform batch BO iterations with a Multi-point Expected Improvement heuristic method with local penalties to obtain the predicted affinity distribution and propose multiple candidate positions in parallel, as shown in Fig. 4b. Finally, optimized sequences were generated by AptaDiff and their binding affinity was evaluated through SPR experiments. This process was iteratively repeated until the binding affinity of the optimized sequences reached or exceeded that of the positive control. We employed the GPyOpt package [65] to implement Bayesian optimization. For the targets IGFBP3 (dataset A) and PTK7 (dataset B), we conducted one round and two rounds of Bayesian optimization, respectively. After generating the final eight new candidates, we achieved approximately 87.9% and 60.2% improvements in binding affinity (the RU value) for the aptamers (A-BO1-0, B-BO2-0, as shown in the Supplementary Table S5, S8 and Fig. 4c, 4e). Fig. 4d and 4f depict the KD values^2^ of the aptamers’ binding to the targets in datasets A and B before and after Bayesian optimization, which also reflects the affinity between the aptamer and the target. It is evident that after Bayesian optimization, the KD values of the obtained aptamers significantly improved. For both target proteins, the minimum KD values for the optimized sequences decreased by 3.6-fold and 2.4-fold, respectively, compared to before optimization (Supplementary Table S4-S8). These results demonstrate that AptaDiff can propose aptamer candidates in an affinity-guided manner and offers the opportunity to optimize their affinity.

Compared to the top 5 affinity aptamers selected through traditional SELEX screening experiments, the aptamers generated after Bayesian optimization exhibit higher overall affinity (see Fig. 5a and 5c, Supplementary Table S10 and S11). Surprisingly, in the experiment targeting the PTK7 protein, among the top 5 generated aptamers with affinity, the 1st-ranked sequence was the same as the 1st-ranked sequence screened by SELEX, and the 4th-ranked sequence was the same as the 2nd-ranked sequence screened by SELEX (sequence B-BO2-0 is the same as B-SELEX-20, B-BO1-5 is the same as B-SELEX-17). In addition, since secondary structure information is crucial for identifying aptamers with affinity, in this section, we used MXFold2 [66] webserver to conduct secondary structure analysis of the nucleic acid aptamers in Fig. 5a and 5c. The results are shown in Fig. 5b and 5d. The top 5 affinity aptamers we generated have similar secondary structures to the top 5 affinity aptamers screened by SELEX, indicating that high-affinity aptamers tend to form similar structures. These findings indicate that our proposed AptaDiff-based aptamer generation pipeline can potentially replace traditional manual screening experiments, generating more high-affinity aptamers, reducing screening time, and contributing to significant savings in manpower, resources, and finances.

**Fig. 5.**
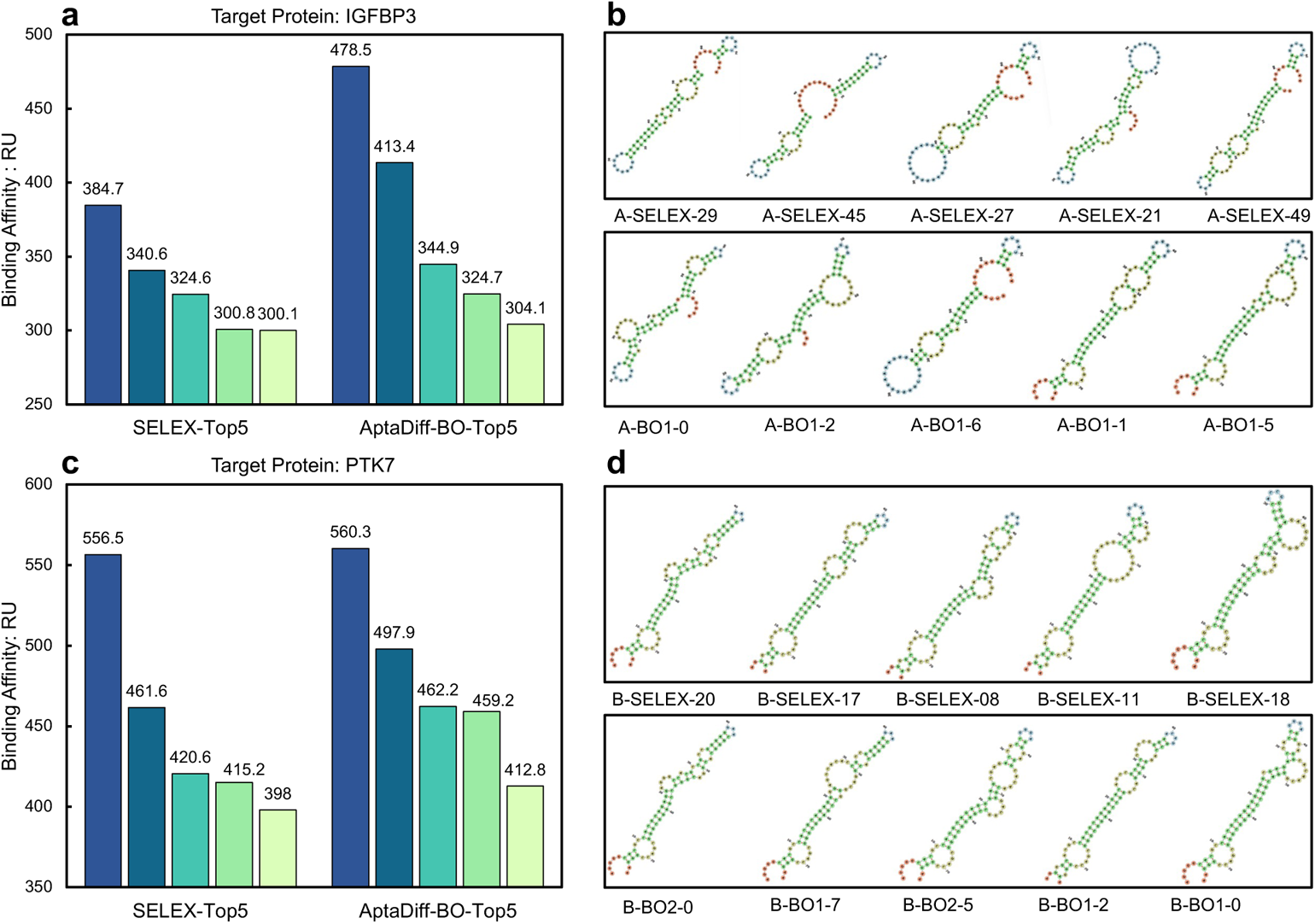
Comparison of the affinity and secondary structure visualization of the top 5 aptamers obtained by SELEX and AptaDiff. **(a)**, Affinity comparison and secondary structure visualization of the top 5 aptamers obtained by SELEX and AptaDiff, respectively, for Dataset A., **(d)** Affinity comparison and secondary structure visualization of the top 5 aptamers obtained by SELEX and AptaDiff, respectively, for Dataset B. The results indicate that, compared to the top 5 aptamers selected through traditional SELEX screening experiments, the aptamers generated after Bayesian optimization by AptaDiff exhibit overall higher affinity.

## Discussion

Generating aptamers that effectively bind to target proteins is significant yet challenging. We introduce AptaDiff, a conditional diffusion model framework capable of generating high-fidelity and novel aptamers. By equipping AptaDiff with a motif-dependent latent space, we enhance interpretability and enable the generation of aptamers with desired binding characteristics through affinity-driven optimization. AptaDiff is the first deep learning framework to demonstrate the power of diffusion generative models on the discrete aptamer sequence space, generating sequences with attributes akin to SELEX-selected aptamers without exploring the challenging sequence space spanning up to 30 − 40 nucleotides.

Presently, aptamer generation methods are broadly categorized into two types: those based on base mutations and those based on deep learning. Hoinka et al.[42] introduced several tools, including AptaCluster, AptaMut, and AptaSim, using mutation information from polymerase error rates for aptamer discovery. Bashir et al. [27] generate new aptamers by randomly mutating 4-nt bases and predict high-affinity aptamers using a trained machine learning model. This method only conducts mutations and evaluates them based on existing sequencing data, and is highly random. Jinho et al. [34] proposed an LSTM-based method using SELEX data but did not consider sequence clustering information. Iwano et al. [39] proposed a VAE-based method using HT-SELEX data, generating sequences not in the sequencing data but with lower fidelity.

In contrast, AptaDiff generates aptamers based on a diffusion model while considering sequence clustering information. It can generate sequences not present in the sequencing data (as shown in Fig. 2a, positive edit distances indicate sequences not present in the original SELEX data). Moreover, we have demonstrated that our generated sequences closely match the original data across multiple attribute metrics, outperforming existing state-of-the-art models on multiple datasets (Fig. 2a, 2b, 3a, and 3b, Table 1). This underscores the higher quality of sequences generated by AptaDiff relative to the aforementioned methods. Consequently, we consider AptaDiff a superior aptamer generation model compared to the methods described above, and we believe it will be a key tool for efficient aptamer discovery. However, AptaDiff does not support end-to-end training and requires a specialized VAE model to obtain a motif-dependent latent space before training the diffusion model. Future research should explore joint training of VAE and diffusion models.

In summary, we have presented a set of open-source conditional discrete diffusion models that provide a foundation for sequence-based nucleic acid aptamer engineering and design. The AptaDiff model can be directly employed for unconditional, affinity-guided aptamer sequence generation. We envision several possible avenues for extending AptaDiff. Firstly, on the model front, considering secondary structure features during the diffusion process would enhance AptaDiff’s performance.

One potential approach in this direction is to use diffusion models to co-design the sequence and secondary structure of aptamers. Secondly, in terms of data, employing aptamer sequences from earlier rounds of HT-SELEX experiments for training data could further expedite the SELEX screening process. Thirdly, in terms of application scenarios, AptaDiff could be used to generate a large pool of aptamer candidates for specific target proteins, forming an aptamer sequence library for SELEX screening, thereby increasing the likelihood of obtaining effective high-affinity aptamers.

As aptamers bind to target proteins through structural complementarity and binding data for aptamer-protein pairs are limited, assessing the binding affinity currently relies on wet-lab experiments, such as SPR. In the future, once a sufficient number of binding data for aptamer-protein pairs accumulates, it may become feasible to design *de novo* aptamers without wet-lab experiments in the loop. Furthermore, in the event of improved computational resources in the future, molecular dynamics simulations can be employed to model the binding conformations between aptamers and target proteins. The accumulation of a substantial amount of simulated data through such simulations could also significantly enhance computational aptamer design. These future prospects hold the potential to revolutionize the field of aptamer design, eliminating the need for extensive wet-lab experimentation and providing a more efficient and versatile approach to aptamer development.

## Data availability

The source data used to train and evaluate our models are available at https://github.com/wz-create/AptaDiff. The HT-SELEX sequences of datasets A and B are available as DRA009383 and DRA009384 in DDBJ.

## Code Availability

The code to reproduce our experiments is available at https://github.com/wz-create/AptaDiff under an MIT License.

## Author Contribution

Z.W. and X.L. conceived the research project. X.L., M.H., and Y.P. supervised and advised the research project. Z.W. designed and implemented the AptaDiff framework. Z.W., Z.L., Y.L., and Y.F. conducted the computational analyses. Z.W., W.Z., L.S., D.H., and L.Z performed the wet experiments and analyses. Z.W., Z.L., Y.L., and X.L. wrote the manuscript. All the authors discussed the experimental results and commented on the manuscript.

## Acknowledgments

We acknowledge helpful discussions with members of the AIM lab. The authors thank the anonymous reviewers for their valuable suggestions.

## Supplementary Figures

**Fig. S1.**
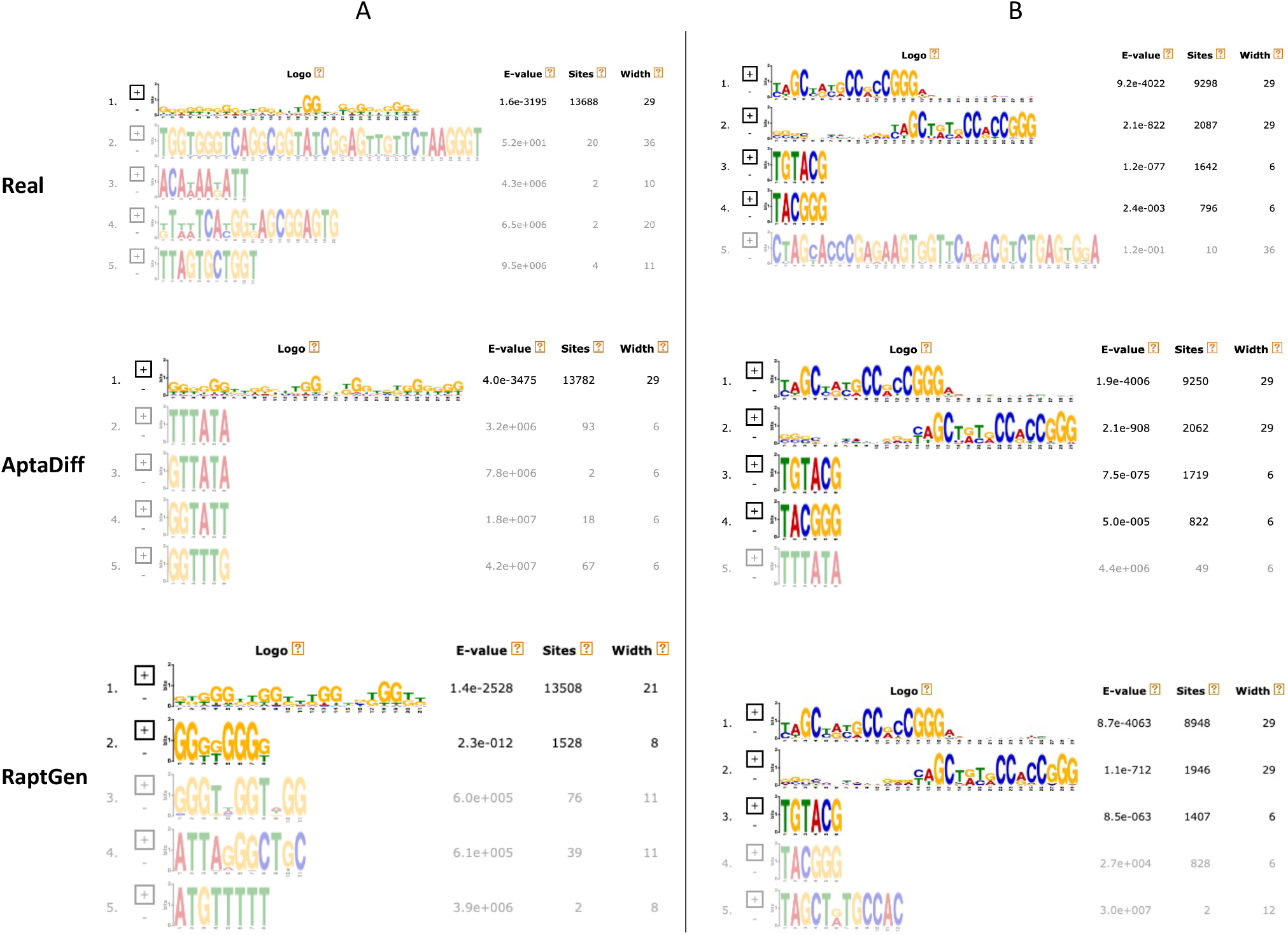
Illustration of the motif clustering analysis by MEME. The left side of the figure presents the results on dataset A, while the right side shows the results on dataset B. ”Real” denotes the real SELEX data, ”AptaDiff” represents data generated by AptaDiff, and ”RaptGen” represents data generated by RaptGen. The E-value indicates the clustering score obtained by MEME, where a smaller score suggests better clustering for motifs in that category. ”Sites” denotes the number of sequences in that category, and ”Width” indicates the maximum length of motifs in that category. The darkness of the sequence logo’s colors is proportional to the “Sites” value, with darker colors indicating a higher number of sequences.

**Fig. S2.**
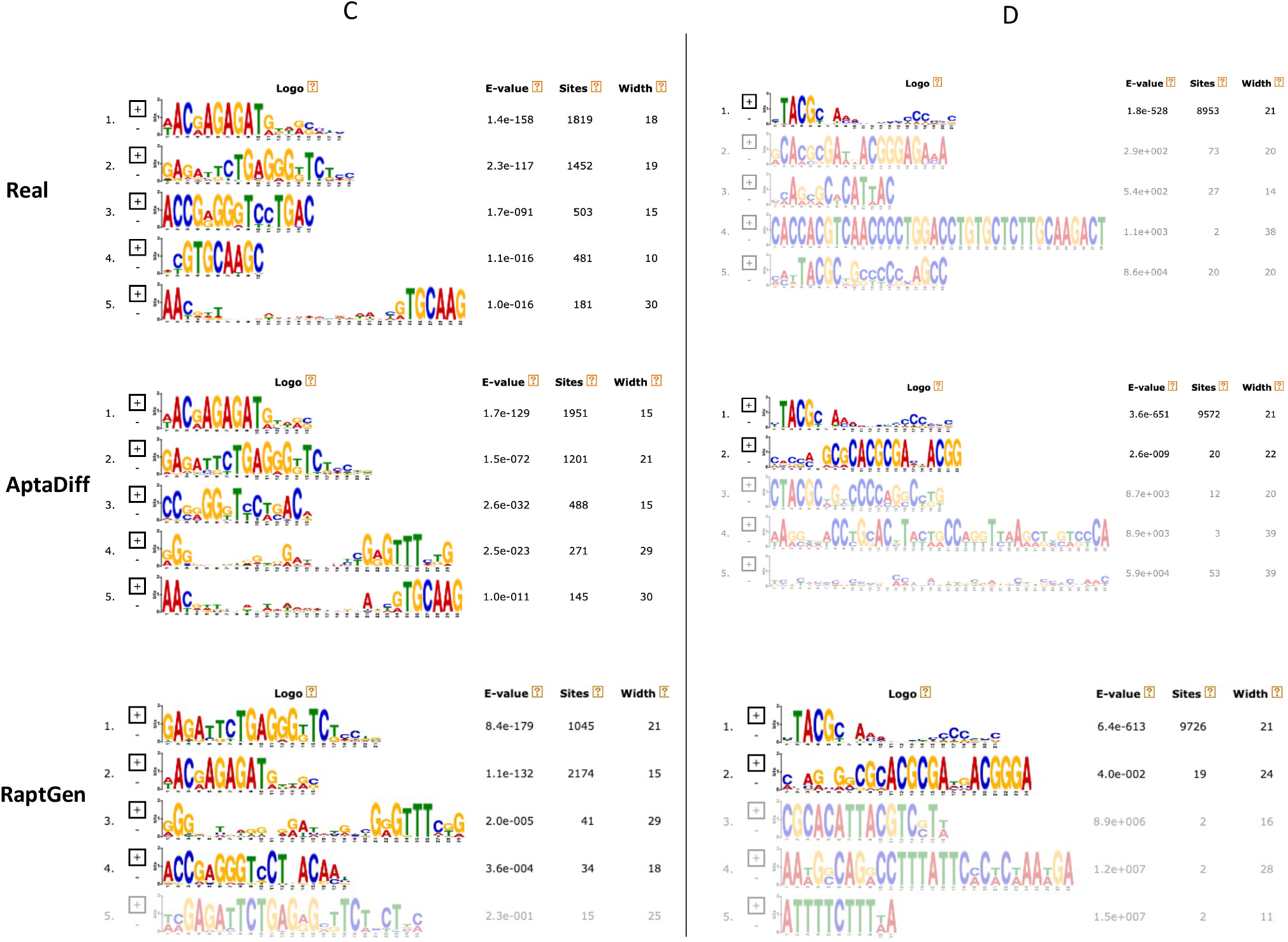
Illustration of the motif clustering analysis by MEME. The left side of the figure presents the results on dataset C, while the right side shows the results on dataset D. ”Real” denotes the real SELEX data, ”AptaDiff” represents data generated by AptaDiff, and ”RaptGen” represents data generated by RaptGen. The E-value indicates the clustering score obtained by MEME, where a smaller score suggests better clustering for motifs in that category. ”Sites” denotes the number of sequences in that category, and ”Width” indicates the maximum length of motifs in that category. The darkness of the sequence logo’s colors is proportional to the ”Sites” value, with darker colors indicating a higher number of sequences.

**Fig. S3.**
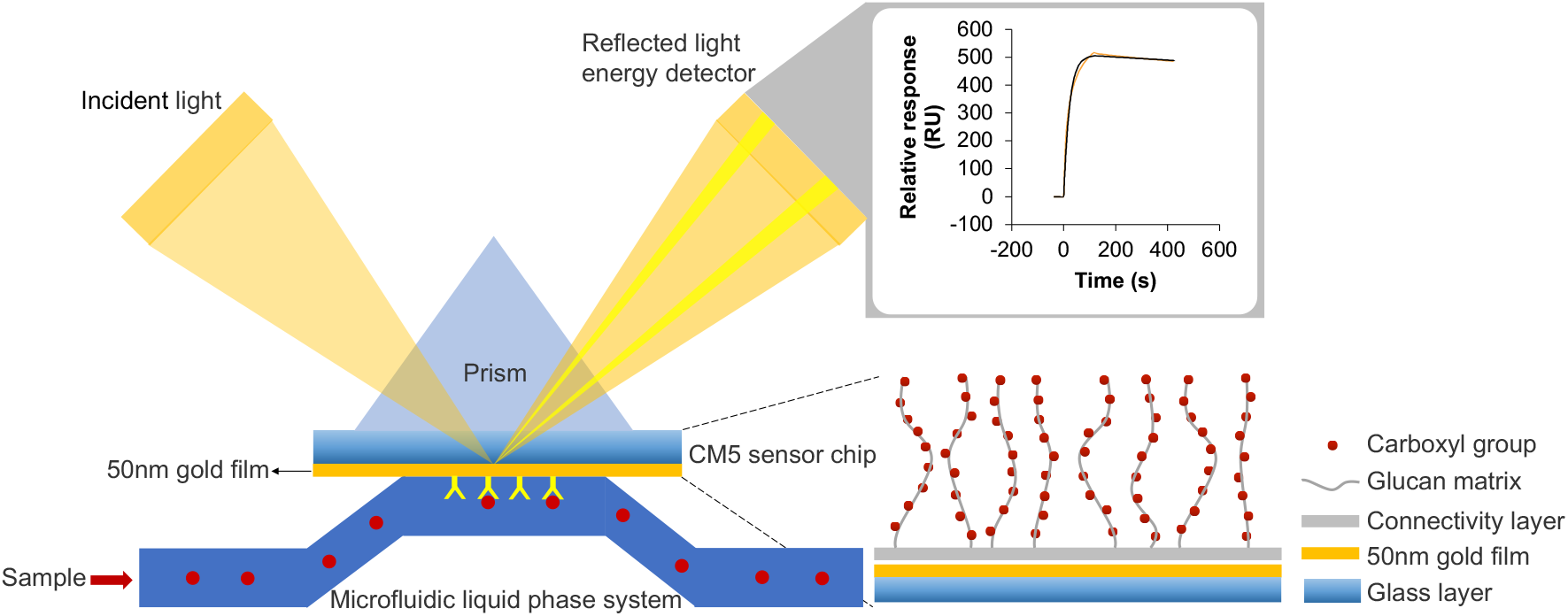
Surface plasmon resonance (SPR) principle. In SPR experiments, when a sample solution (e.g., aptamers) flows over the surface of a chip coated with biomolecules (e.g., antibodies), interactions between molecules in the sample and those on the surface result in an increase in surface mass, consequently causing a change in the resonance angle frequency. RU values are employed to quantify the extent of this change in surface mass. Typically, 1 RU signifies the change in resonance angle frequency generated when biomolecules with a mass of 1 nanogram (ng) per square millimeter (mm^2^) bind to the surface.

**Fig. S4.**
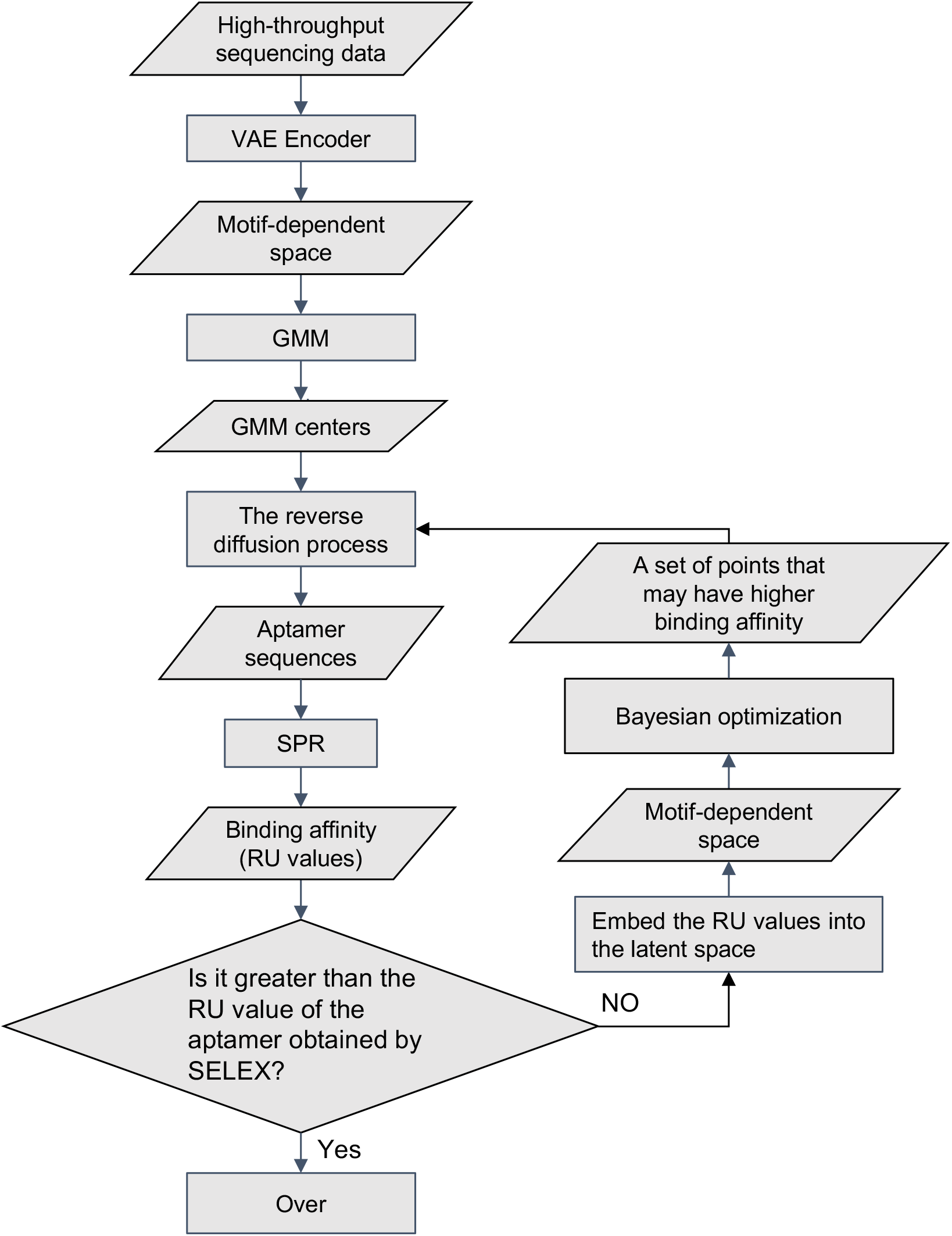
Flowchart of the actual application of trained AptaDiff. Upon completing AptaDiff training, we employ a Gaussian Mixture Model (GMM) to classify points within the latent space, subsequently selecting the category center points of GMM as starting conditions to generate sequences through the reverse diffusion process. These sequences then undergo SPR experiments to measure their binding affinity (i.e., RU values) with target proteins. These RU values are subsequently embedded back into the latent space, after which Bayesian optimization is applied to identify a set of points likely to have higher binding affinity. These points, in turn, serve as conditions to generate sequences through the reverse diffusion process for further SPR experiments. This cycle continues until the candidate sequences’ affinity surpasses those identified through traditional SELEX experiments.

## Supplementary Tables

**Table S1.**
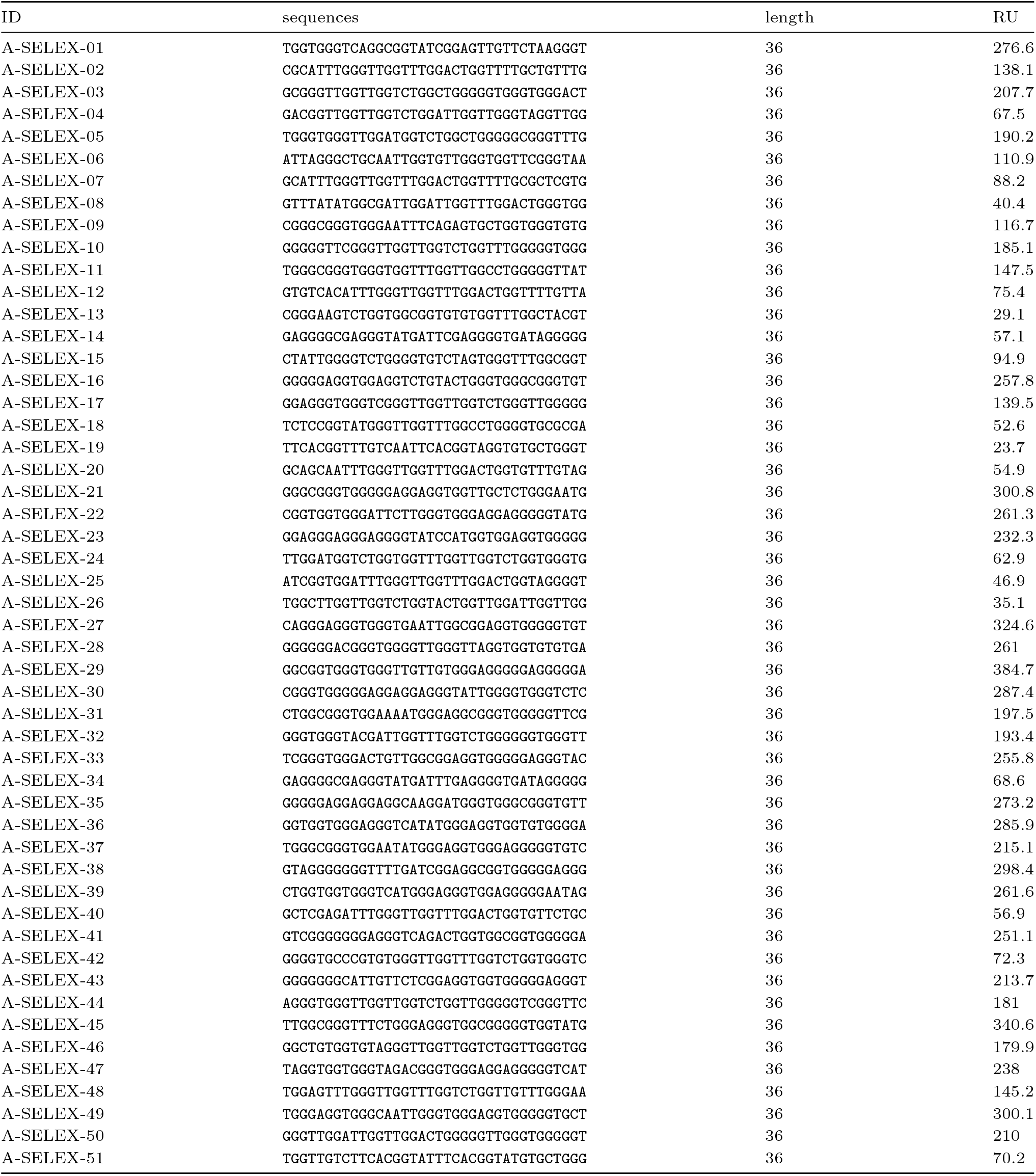
Sequences hand-picked in SELEX experiments targeting the IGFBP3 protein. The aptamer sample concentration in the SPR experiment was 500 nM.

**Table S2.**
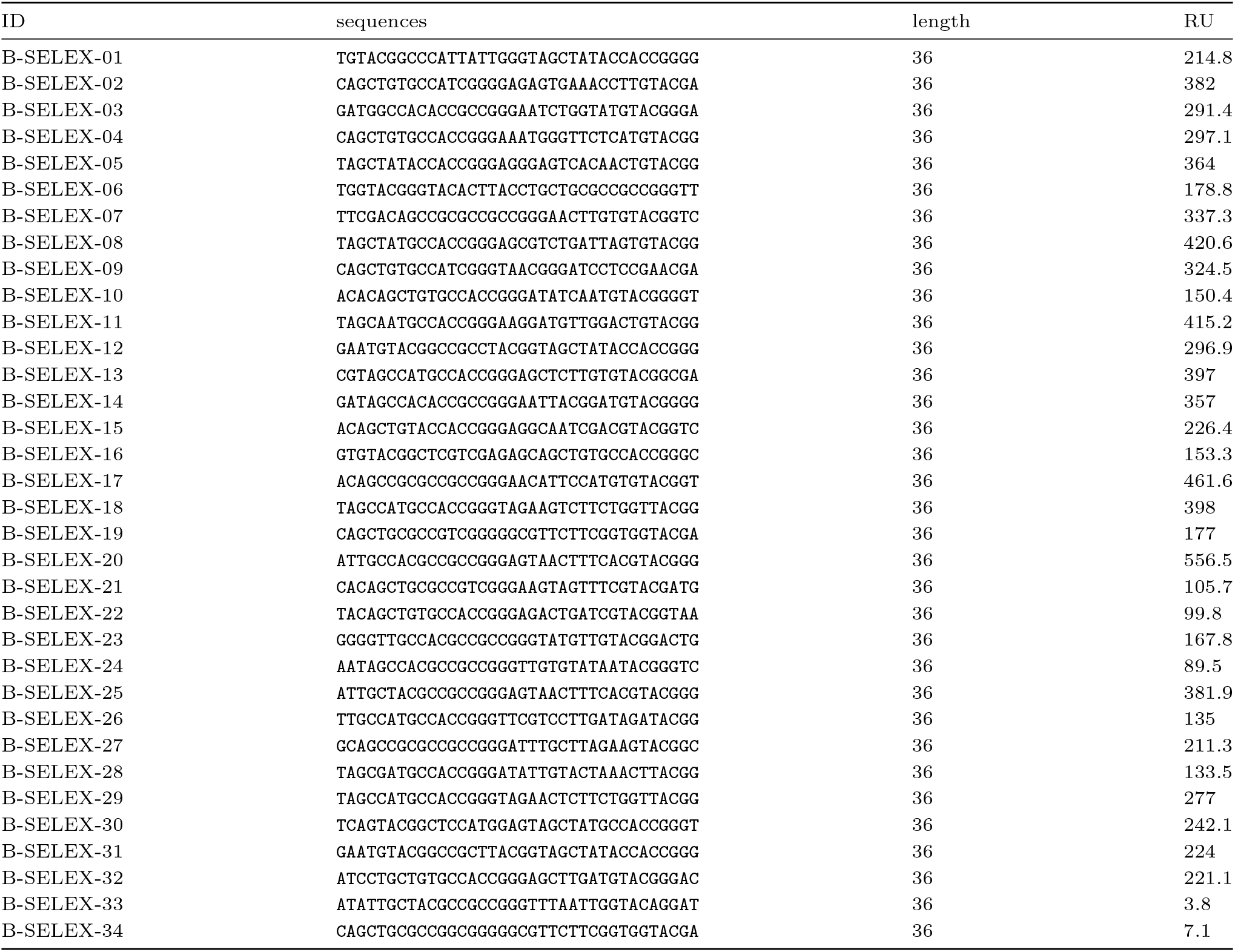
Sequences hand-picked in SELEX experiments targeting the PTK7 protein. The aptamer sample concentration in the SPR experiment was 500 nM.

**Table S3.** Quantitative Illustration of the generative performance on datasets A, B, C, and D. FID reported on 10k samples (Lower is better). Using DNA-BERT2 [59] as the feature extractor.

| Source | Dataset | Target Protein | FID |  |
| --- | --- | --- | --- | --- |
|  |  |  | RaptGen | AptaDiff |
| Ours | A | IGFBP3 | 1.2627 | 0.0222 |
|  | B | PTK7 | 0.0281 | 0.0067 |
| Public | C | TGM2 | 0.0293 | 0.0097 |
| | D | ITGαVβ3 | 0.0233 | 0.0137 |

## Supplementary Sections

### Supplementary Section 1: Sequences Obtained in the Present Study

The sequences obtained in the present study are shown in Supplementary Table S4-S8. The ID is named after a rule of dataID, the method to select the sequence, and the index of the sequence. The dim1 and dim2 represent the two dimensions of the two-dimensional latent space. RU and KD reflect the binding affinity of the target sequence as assessed by the SPR experiment. The sample concentration in the SPR experiment was 500 nM.

### Supplementary Section 2: Statistics of the Dataset

The SELEX experiments for datasets A and B were conducted against target proteins IGFBP3 and PTK7, respectively. For each protein target, six rounds of SELEX screening were carried out, with each round encompassing the following steps: (1) positive selection where the ssDNA library was exposed to the target protein, (2) elution of nucleic acid molecules bound to the target protein followed by PCR amplification and enrichment of the library, and (3) high-throughput sequencing of the enriched library. Considering that candidate aptamer libraries enriched in later rounds of the SELEX experiments have a higher degree of enrichment, we selected the data obtained from the last round of experiments as the initial data for datasets A and B. Following this, we processed the data to get the datasets for model training. Specifically, all sequences in the training set were carefully filtered. Sequences meeting the following three criteria were retained: (1) exact matching with primers (including forward and reverse primers), (2) precise alignment with the sequence design length, and (3) sequences read more than once. The statistics of the datasets are shown in Table S9. The column names are as follows:

- Source: Source of the datasets, where “Ours” indicates that the dataset comes from our own SELEX experiment, “Public” represents the public SELEX dataset obtained from DDBJ [50, 51].
- Target: The target protein.
- ID: The ID named after the rule; indicator of the dataset-round of the SELEX.
- Filtered unique: Number of unique sequences matched by primer match and target length.
- Target length: The designable target length is the length of the variable random region in the middle of the aptamer.

Note that datasets C and D are taken directly from the real training data used in Reference [39].

### Supplementary Section 3

Comparative Analysis to Traditional SELEX Screening Experiments.

Compared with the top 5 affinity aptamers selected by traditional SELEX screening experiments, the overall affinity of the aptamers we generated after Bayesian optimization was higher (Table S10 and S11)

### Supplementary Section 4

Details of Handling the Diffusion Model Stochasticity.

During the sample generation process following the training of AptaDiff, two main determinants influence the outcome: the latent representation (**z**) from the VAE and the random noise point (***x***_*T*_) sampled from a multinomial distribution. Our design leverages the latent representation to steer the diffusion generation process. To secure the determinacy and reproducibility of the samples generated by AptaDiff, thereby ensuring that the generated samples are influenced solely by the latent representation, we adhere to the approach suggested in reference [53]. Specifically, we assign a fixed random seed to each latent representation (**z**) to sample the nosie (***x***_*T*_) and share this stochasticity during the reverse diffusion process across all generated samples. By adopting this methodology, we ensure that each generated sample can be deterministically associated with a given latent representation, allowing for the reproducibility of results which is crucial for the verifiability and scientific rigor of our findings.

**Table S4.**
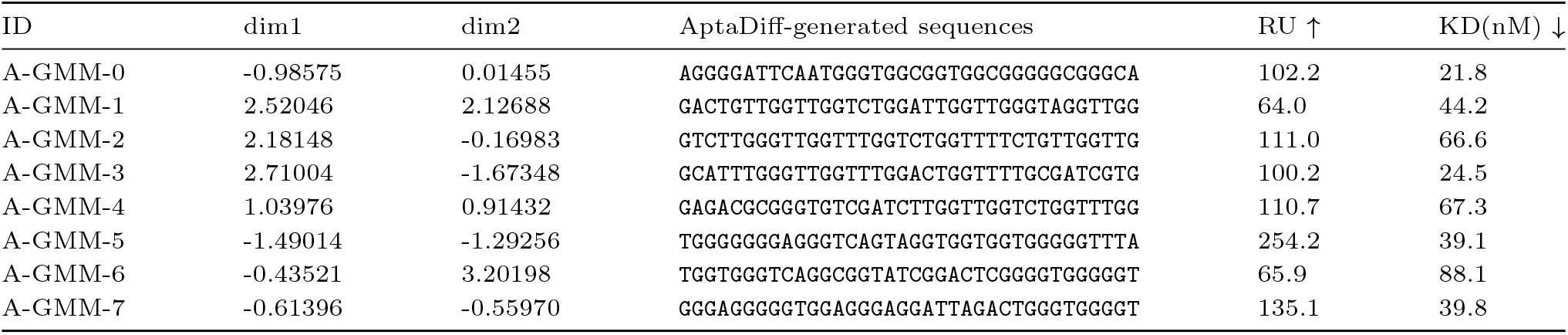
The sequences obtained in the initial sequence selection for dataset A.

**Table S5.**
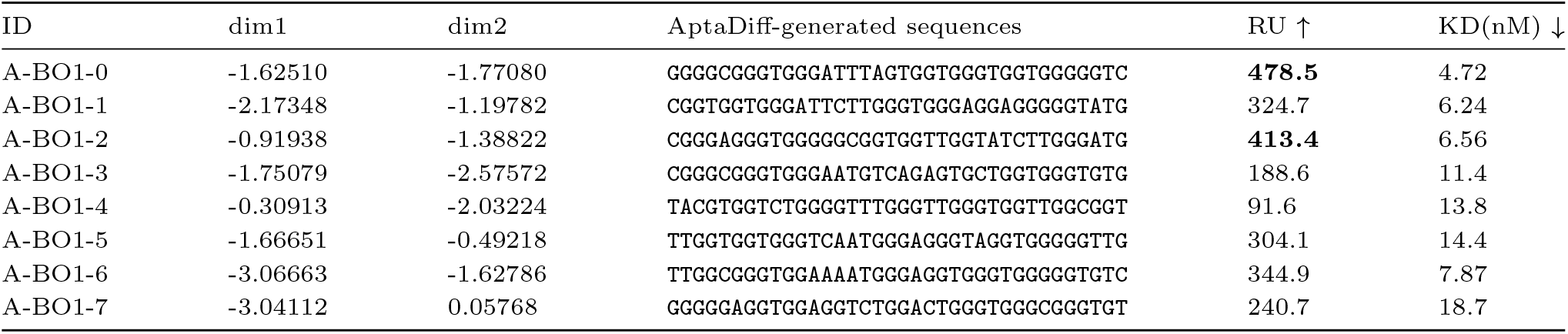
The sequences obtained after the first Bayesian optimization for dataset A.

**Table S6.**
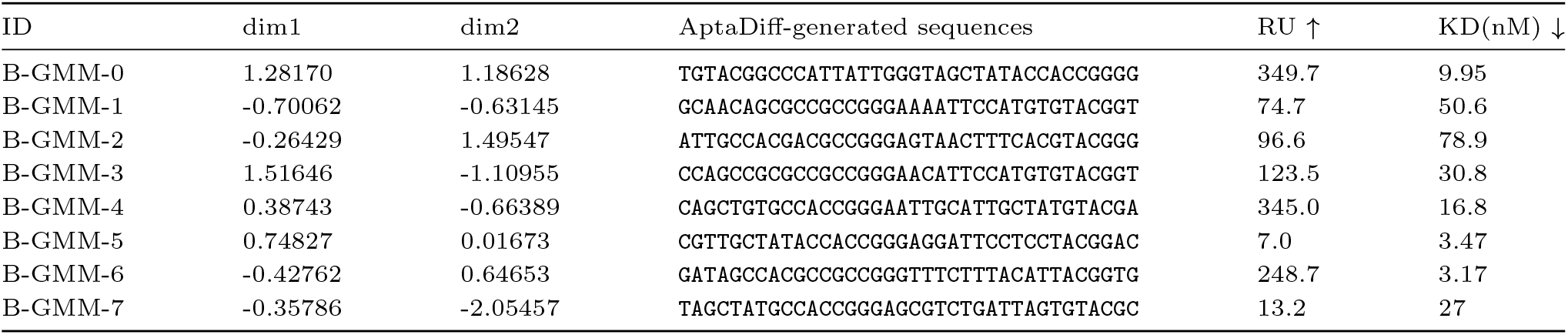
The sequences obtained in the initial sequence selection for dataset B.

**Table S7.**
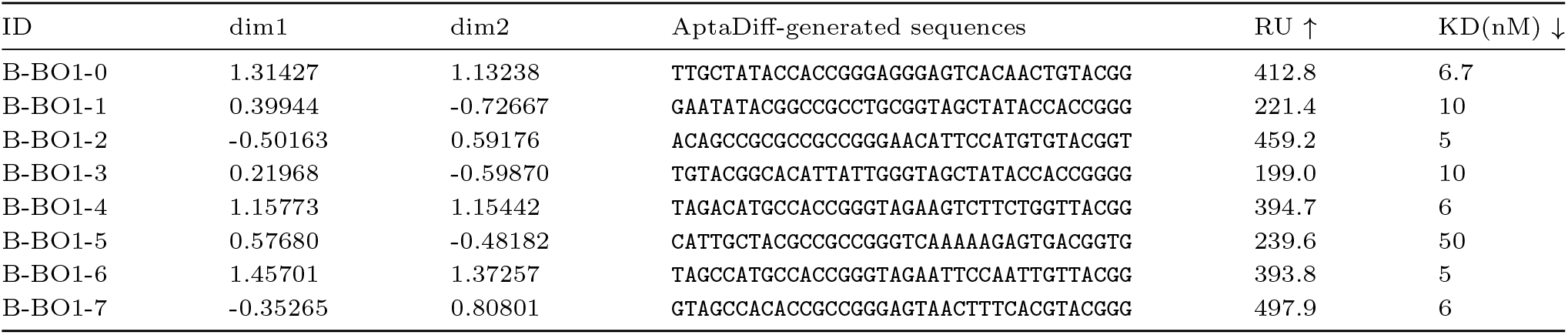
The sequences obtained after the first Bayesian optimization for dataset B.

**Table S8.**
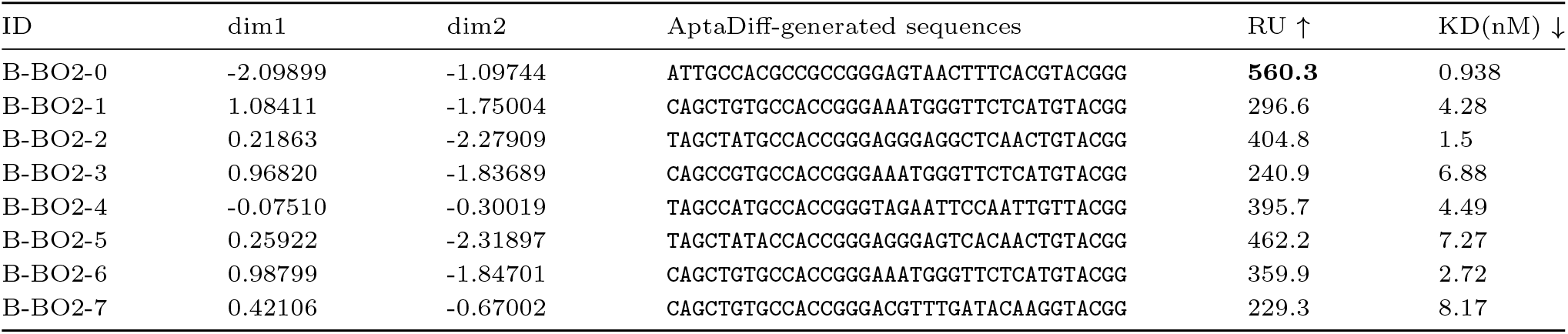
The sequences obtained after the second Bayesian optimization for dataset B.

**Table S9.** Statistics of the datasets used in our research.

| Source | Dataset | Target | ID | Filtered unique | Target length |
| --- | --- | --- | --- | --- | --- |
| Ours | A | IGFBP3 | A-6R | 13802 | 36nt |
|  | B | PTK7 | B-6R | 11532 | 36nt |
| Public | C | TGM2 | C-4R | 38613 | 30nt |
| | D | ITGαVβ3 | D-4R | 31358 | 40nt |

**Table S10.**
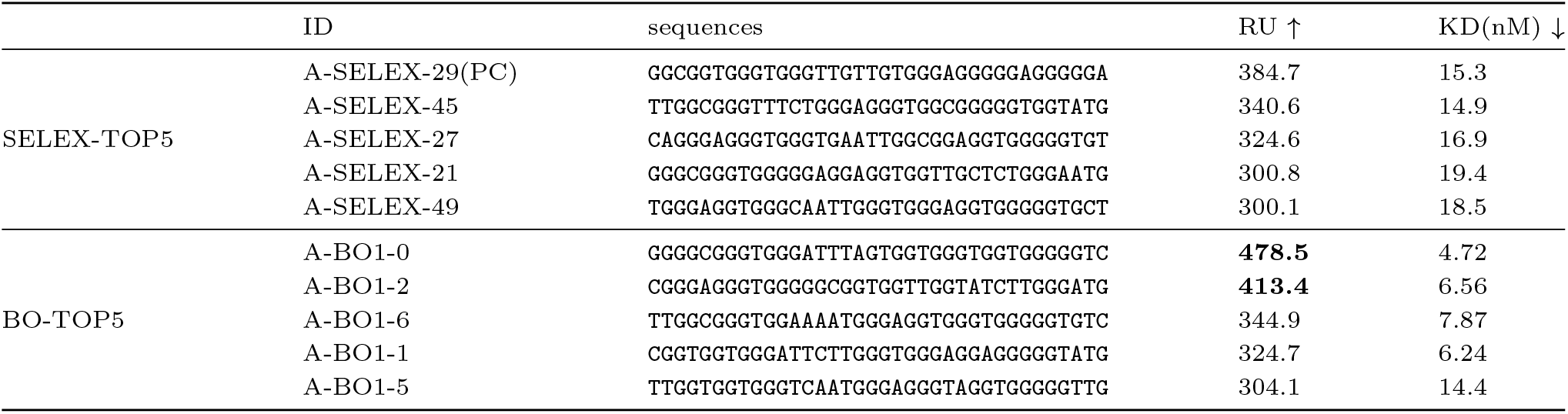
The sequences of the top 5 affinity aptamers we generated and the top 5 affinity aptamers selected through traditional SELEX screening for dataset A.

**Table S11.**
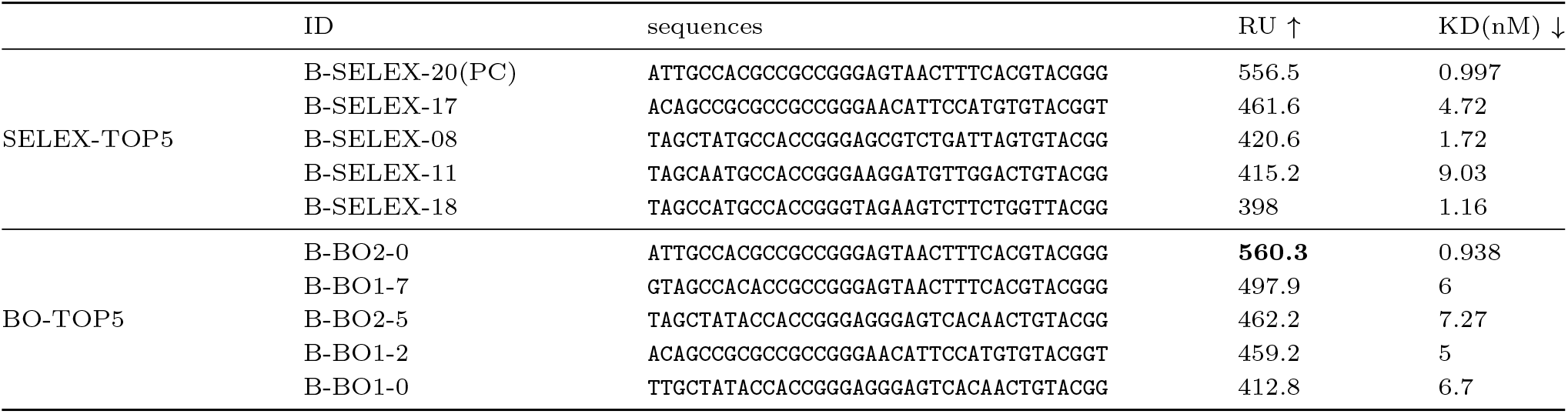
The sequences of the top 5 affinity aptamers we generated and the top 5 affinity aptamers selected through traditional SELEX screening for dataset B.

## Footnotes

1 RU values: Response Units, a measure of the change in refractive index on the sensor surface in SPR experiments, indicating the binding of the molecule to the target. Generally speaking, higher RU values indicate higher affinity.

2 KD value: Dissociation constants, the ratio of the protein sequence dissociation rate constant Kd to the protein sequence association rate constant Ka, a measure of the strength of binding between molecules in a biomolecular interaction. Generally speaking, lower KD values indicate higher affinity.

